# Nematode histone H2A variant evolution reveals diverse histories of retention and loss and evidence for conserved core-like variants

**DOI:** 10.1101/2022.03.02.482035

**Authors:** Swadha Singh, Diana Chu, Scott Roy

## Abstract

Histone variants are paralogs that replace canonical histones in nucleosomes, often imparting novel functions. Despite their importance, how histone variants arise and evolve is poorly understood. Reconstruction of histone protein evolution is challenging due to high amino acid conservation and large differences in evolutionary rates across gene lineages and sites. Here we combined amino acid sequences and intron position data from 108 nematode genomes to trace the evolutionary histories of the three H2A variants found in *Caenorhabditis elegans*: the ancient H2A.Z^HTZ-1^, the sperm-specific HTAS-1, and HIS-35, which differs from canonical H2A by a single glycine-to-alanine C-terminal change. We find disparate evolutionary histories. Although the H2A.Z^HTZ-1^ protein is highly conserved, its gene exhibits recurrent intron gain and loss. This pattern suggests that it is intron presence, rather than specific intron sequences or positions, that may be important to H2A.Z functionality. In contrast, for HTAS-1 and HIS-35, we find variant-specific intron positions that are conserved across species. HIS-35 arose in the ancestor of *Caenorhabditis* and its sister group, including the genus *Diploscapter*, while the sperm-specific variant HTAS-1 arose more recently in the ancestor of a subset of *Caenorhabditis* species. HIS-35 exhibits gene retention in some descendent lineages but also recurrent gene loss in others, suggesting that histone variant use or functionality is highly flexible in this case. We also find that the single amino acid differentiating HIS-35 from core H2A is ancestral and common across canonical *Caenorhabditis* H2A sequences and identify one nematode species that bear identical HIS-35 and canonical H2A proteins, findings that are not predicted from the hypothesis that HIS-35 has a distinct function. Instead, we speculate that HIS-35 enables H2A expression across the cell cycle or in distinct tissues; genes encoding such partially-redundant functions may be advantageous yet relatively replaceable over evolutionary times, consistent with the patchwork pattern of retention and loss of both genes. Our study shows the evolutionary trajectory for histone H2A variants with distinct functions and the utility of intron positions for reconstructing the evolutionary history of gene families, particularly those undergoing idiosyncratic sequence evolution.

## INTRODUCTION

All characterized eukaryotic cells compact their DNA into nucleosomes, which consist of an octameric histone complex comprised of two copies each of the core histone proteins H2A, H2B, H3, and H4, and which wrap around ∼147 bps of DNA (1-4). In many species, genes encoding canonical core histones (also referred to as ‘replication-dependent histones’) do not contain introns and are organized into multiple gene copies and in a tandem-repeat structure, which facilitates their rapid, coordinated expression (4-7). The expression of canonical histones is tightly coupled with the S-phase of the cell cycle because of the critical need for the large bulk of histone proteins required to package and compact the newly synthesized DNA (8-10).

Histones have evolved variant forms that further regulate chromatin compaction and affect processes like transcription, DNA repair, and development (11-20). Such histone variants are often considered to be a part of the ‘histone code’ and control distinct sets of genes during specific times and tissues in both animals and plants (11, 12, 21-23). Among all the core histones, H2A is the fastest evolving histone, showing the most diversity in the variant types and expression patterns (24, 25). For example, the highly conserved H2A.Z is present in every eukaryotic species and ubiquitously expressed. It has been shown in different species to play a variety of roles including transcriptional regulation, heterochromatin boundaries, DNA repair, DNA replication, and dosage compensation (26-31). In contrast, H2A.X has arisen independently in several lineages, with functions in DNA damage response and transcription (18, 31, 32), while macroH2A, which is involved in × inactivation and stress response, is present only in some lineages, making its origins unclear (3, 4, 24, 33-36). Finally, short histone variants, such as H2A.B, H2A.P, H2A.L, and H2A.Q, are expressed in testis (except H2A.B which is also expressed in the brain cells) (25, 37-40). In short, a great wealth of histone H2A variants are observed, ranging from those shared across all eukaryotes to others that are species-specific (3, 4, 18, 19, 35). H2A variants also have distinct functions in a broad range of processes and can be ubiquitously expressed or expressed only in certain tissues (11, 25, 41, 42). However, with the exception of the well-studied ancient variant H2A.Z, how histone H2A variants have arisen and evolved remains unknown.

Clues to how histone variants differ may stem from the distinct gene structures and expression patterns compared to canonical histones. Unlike canonical histone genes, variant histone gene expression is not restricted to S phase expression and can be tissue-specific. Most histone variants are typically found in a single copy in the genome and contain introns in their pre-mRNAs (5-7, 43, 44). Notably, the roles of these introns in variant function remain poorly studied, including whether the introns themselves impart novel functionality. For instance, sequences found within introns of other genes can regulate gene expression (43, 45-48). Introns can also allow for alternative splicing, which is common in animals and plays important role in development and disease (49-52). Introns have been used in other studies to determine the evolutionary trajectory of other protein families (53-57). Thus, the presence of introns in histone variants may provide useful clues to exploring the histone evolution and function across closely related species.

Core histone functions are expected to be highly conserved across eukaryotes, given their central roles in ensuring DNA packaging and protection (4, 61). Thus, observed protein changes in core histones are expected to be largely neutral with respect to protein function. On the other hand, amino acid differences between variant histones and core histones are thought to generally lead to functional differences (3, 4, 62). Indeed, protein sequence differences between well-studied variants and their core homologs have been shown to affect chromatin structure and function. The histone H3.3 variants, for example, which differ from each other by 4-5 amino acids, play distinct roles in transcriptional activation, chromatin remodeling, heterochromatin formation, and development (14, 16, 63-66). On the other hand, the functional significance of protein differences between variant histones and their core counterparts in some cases is unclear. For instance, the protein sequence of *C. elegans* HIS-35 differs by a single ‘A’ instead of a ‘G’ at the 124th position from the S-phase *C. elegans* H2A, the functional significance of which is unknown.

In this study, we focus on the evolutionary histories of the three very different H2A variants found in *Caenorhabditis elegans:* H2A.Z^HTZ-1^, HIS-35, and HTAS-1 (Figure 1). H2A.Z^HTZ-1^ is an ortholog of the evolutionarily conserved variant, H2A.Z. HIS-35 intriguingly differs from core H2A by a single amino acid difference. HTAS-1 shows a greater divergence from core H2A, particularly at the highly divergent C and N termini, and appears to be expressed only in sperm and has only been reported in *C. elegans* to date (67). The presence of a discrete number of H2A variants with distinct functions and/or expression patterns in combination with the availability of sequence data across a large number of nematode species allows a unique opportunity to track the evolutionary trajectory of this histone variant family.

**Figure 1.**
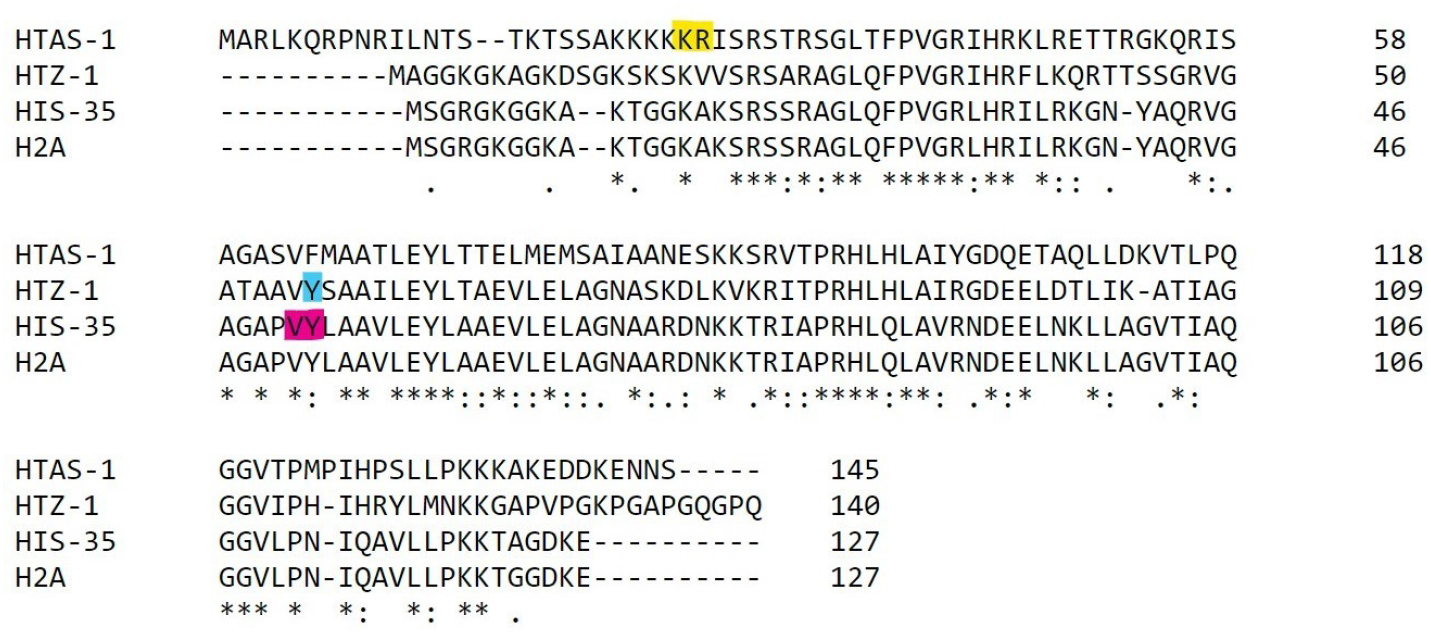
Sequence and intron position comparison of three H2A variants of *C. elegans. C. elegans* core histone H2A and its three variant paralogs contain different sequences and intron positions. The variants differ either in intron positions or the phases in which they interrupt the codon. HTAS-1 (highlighted with a yellow) has a phase 0 intron between the 26th and 27th amino acid; HIS-35 (highlighted with pink) has a phase 0 intron between 50th and 51st amino acid. HTZ-1 (highlighted with blue) has a phase 2 intron splitting the 57th amino acid.

## RESULTS

### Sequence-based phylogenetic methods fail to reconstruct H2A gene family evolution

Taking advantage of vast sequencing (68-70) efforts across nematode species, we used BLAST searches across 108 reported nematode genomes to identify all annotated copies of H2A and H2A-related gene variants across the diversity of nematodes. After filtering and collapsing identical proteins, we were left with 408 unique sequences. We then used standard phylogenetic methods to reconstruct the evolutionary history of these sequences (Figure S1, intronless (thus putative core or canonical) H2As are highlighted in yellow, HTZ-1 homologs are highlighted in pink, putative HIS-35 homologs are highlighted in blue, and putative HTAS-1 homologs are highlighted in gray). However, scrutiny of the recovered phylogenetic tree revealed several bizarre findings. For instance, core H2A proteins were grouped as separate clades that included very deeply-diverged nematode sequences; on the other hand, species-specific variants were often found grouped far from core proteins from the same or related species. Perhaps the clearest case can be shown by performing phylogenetic reconstruction on core H2A orthologs and on the HIS-35 orthologs identified based on intron positions (described in detail below). Here, we expect distinct clades of H2A and HIS-35 sequences, yet we recover no such separate clades (Figure S2, HIS-35 is highlighted).

Some of these anomalies are as expected by errors in phylogenetic reconstruction due to model misspecification (the phenomenon in which differences between the assumed model of sequence evolution and the actual evolutionary process led to errors in phylogenetic reconstruction) (71). The possibility of model misspecification is increased in the case of histone genes by several factors: extreme differences in rates across sites (some sites show conservation across eukaryotes, others are variable within *Caenorhabditis*); large relative differences in rates across gene lineages (the extremely slow evolution of core proteins, but substantially more change in some variants); and the generally small number of total sites given the short length of histone proteins.

### Intron position and phase distinguish the three H2A variants of *C. elegans*

As an alternative approach, we used another potential source of phylogenetic information: the position of the spliceosomal introns that disrupt nuclear genes, including variant histones. Intron positions can be conserved over very long times in orthologous genes (53-56, 58, 59). Whereas early work considered the possibility of intron ‘sliding’, in which an intron would migrate a few base pairs along a gene, recent work has shown that intron sliding is a very rare occurrence, and thus that intron positions are very often conserved over very long times (54, 59). For example, in *Theileria* apicomplexans, 99.7% of intron positions are conserved between *T. parva* and *T. annulata*, diverging roughly 82 million years ago (53). In mammals, 99.9% of intron positions are conserved between humans and dogs, diverging around 100 million years ago (55).

We began our study of intron-exon structures in H2A variants by obtaining intron-exon structures for all H2A gene family members and performing alignments to determine intron position sharing across genes. We first aligned the three main *C. elegans* H2A variants (Figure 1), HIS-35, HTAS-1, and H2A.Z^HTZ-1^, each of which has a single intron position (highlighted boxes in Figure 1). Scrutiny of the intron positions showed that the three genes contain introns at different positions, differing in the codon that they interrupt or and/or in the phase at which they disrupt the codon. Interestingly, the intron position in HIS-35 falls very near to that found in H2A.Z^HTZ-1^. However, these introns are unlikely to represent a shared intron, for two reasons. First, intron positions rarely slide between phases (54, 59). Second, HIS-35’s near identity to H2A (see below) strongly suggests that it more likely evolved from intronless H2A by gene duplication and intron gain, and not from H2A.Z^HTZ-1^. Thus, the differences in the intron positions between the three *C. elegans* variants suggest that the three variants evolved independently as different duplicates of core H2A, rather than from each other.

### Intron position and sequence evidence indicates the origin of HTAS-1 within *Caenorhabditis* and subsequent retention and loss

The *C. elegans* sperm-specific H2A variant, HTAS-1, contains a single intron between codons 26 and 27 (“phase 0”; Figure 1, highlighted with yellow). Alignment across all the H2A variants of 108 nematodes revealed 16 genes that share an intron at the exact homologous position as *C. elegans* HTAS-1 (Figure 2) from species falling within a single subclade of 22 species (represented by the red hash mark on the tree branch in Figure 2) within the *Caenorhabditis* genus, suggesting an origin of this intron position within the common ancestor of these species. Scrutiny of the sequence gene tree (Figure S1, highlighted in gray, IP_152) revealed that this same set of genes (i.e., those sharing the intron) appears together as a clade, reinforcing the notion that the genes sharing the HTAS-1 intron position represent a set of orthologs. We next sought to identify potential HTAS-1 orthologs in additional *Caenorhabditis* species, both the 6/22 within the HTAS-1-containing subclade that lack intron-containing HTAS-1 candidates (marked by purple hash on the tree branch in Figure 2) as well as the 10 species outside this subclade. No genes from any such species were found grouping with the putative HTAS-1 ortholog clade; thus, overall, we couldn’t find any other genes that are candidates for HTAS-1 orthologs. In total, the data is consistent with the origin of HTAS-1 and its gene-specific intron occurring in the same ancestor of a subset of *Caenorhabditis* species.

**Figure 2:**
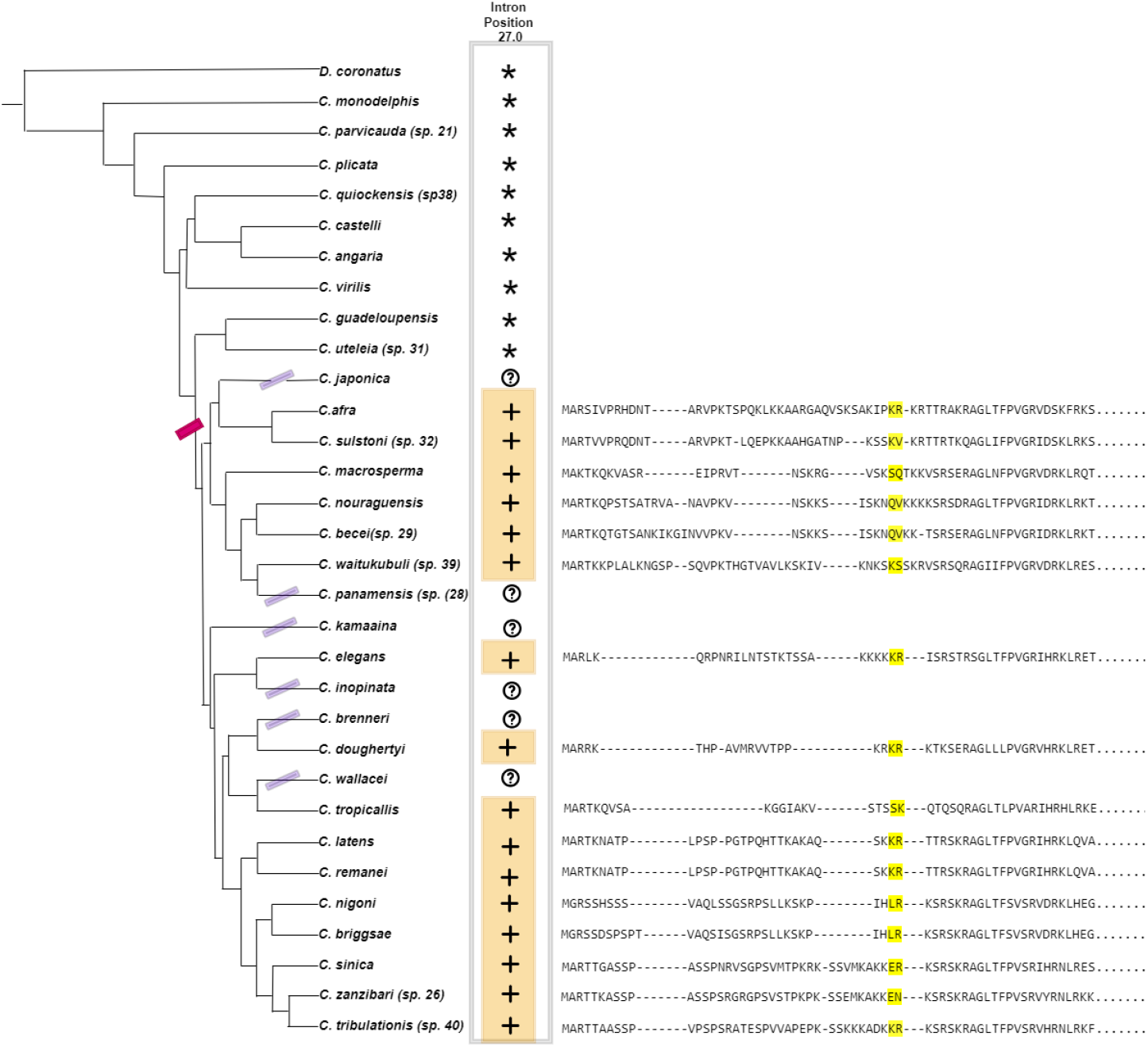
Intron position and sequence evidence indicate the origin of HTAS-1 within *Caenorhabditis* and subsequent loss. On the left is the previously reconstructed species tree of *Caenorhabditis* and *Diploscapter*. The likely origin of HTAS-1 is indicated with a dark pink bar. HTAS-1 presence in a species is indicated by the plus sign. The question mark denotes species where HTAS-1 homologs could not be identified.. The species where HTAS-1 was not incorporated are indicated as an asterisk. On the right is the multiple sequence alignment of HTAS-1 proteins showing the aligned intron positions (highlighted with yellow). For consistency, we show the portion of the phylogenetic tree containing the *Caenorhabditis* species and *Disploscapter*.

As noted above, in six species within the HTAS-1-containing clade of *Caenorhabditis* (*C. wallacei, C. brenneri* and *C*.*inopinata* (*sp34*), *C. kamaaina, C. panamensis* (*sp28*), *and C. japonica*), we could not find annotated genes with either an intron position at the HTAS-1-specific position (Figure 2, marked with purple hashes on the tree branch) or other genes that grouped with the putative HTAS-1 orthologs (Figure S1). However, we were unable to distinguish whether these absences represent true biological loss or failure of gene annotation, which is particularly possible given that histone genes’ short lengths increase the possibility they would be missed by standard annotation methods. Indeed, we identified one species harboring a likely HTAS-1 sequence for which only the second exon was annotated (Figure S1), however, TBLASTN searches against the genome failed to confidently detect this gene. This is likely because, given the much higher rate of evolution of HTAS-1, HTAS-1 from one species often shows a greater degree of core H2A from another species than to its HTAS-1 ortholog; consequently, TBLASTN searches using HTAS-1 from one species typically give many hits, with the likely HTAS-1 not being within the top dozen hits, making it difficult to distinguish gene absence from the failure of annotation. In total, then, the data is consistent with a single origin of HTAS-1 and its characteristic intron position within the ancestor of a subset of studied *Caenorhabditis* species, but we cannot determine with certainty whether HTAS-1 has been lost in up to six independent lineages (Figure 2).

### Intron position conservation suggests HIS-35 originated in the *Caenorhabditis-Diploscapter* ancestor and has undergone subsequent gene retention and loss

The variant HIS-35 of *C. elegans* has a phase zero intron between the 50th and the 51st codon (Figure 1). Alignment across all H2A variants revealed 20 species with genes that share this intron position (Figure 3, marked with a plus sign). In all cases, these genes show very high sequence similarity with H2A at the protein level. These genes are from species falling in the clades of *Caenorhabditis* and its sister genus *Diploscapter*, representing 20/32 species within this clade, suggesting an origin of this intron position within the common ancestor of *Diploscapter* and *Caenorhabditis* (Figure 3).

**Figure 3:**
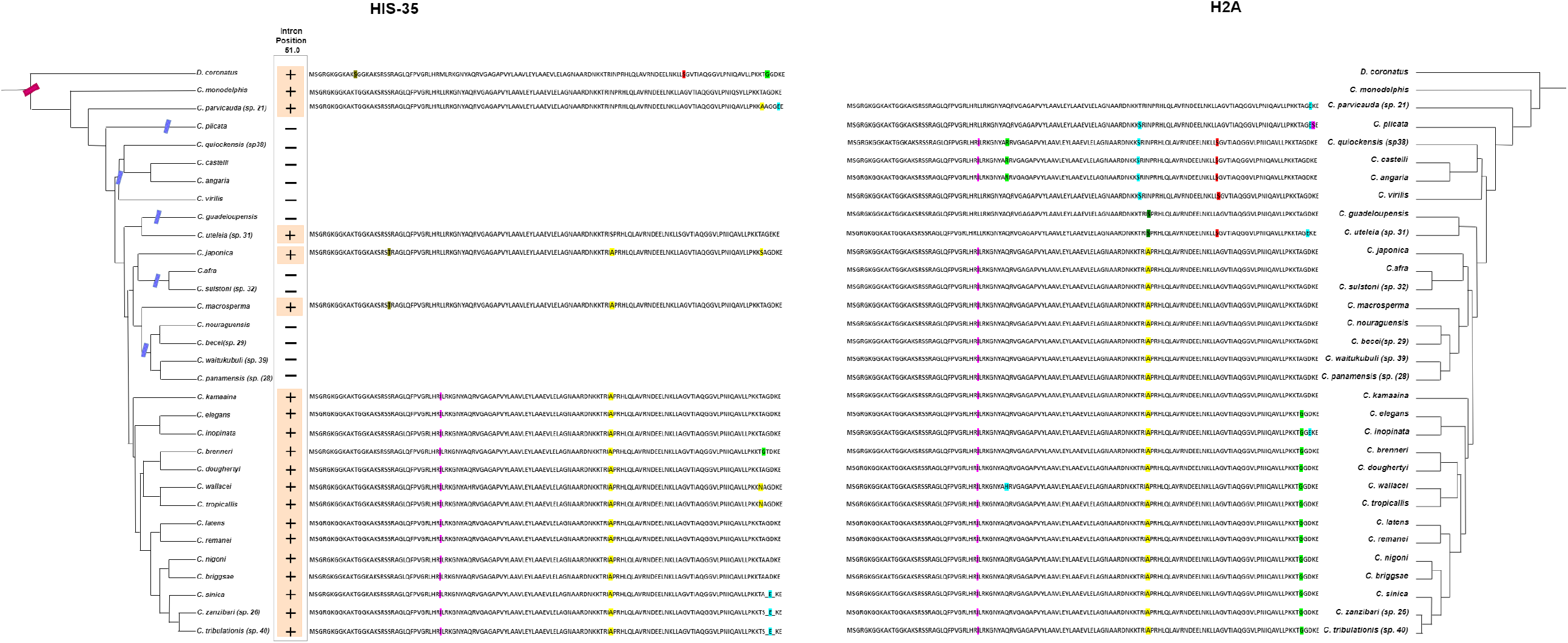
Comparison of the variant HIS-35 (left) with canonical H2A (right) sequences across *Caenorhabditis* species. Inferred sequence changes relative to the reconstructed ancestral sequence are highlighted with different colors (pink, red, yellow, blue, green), with identical changes colored identical colors across the figure (i.e., the L-to-R change at position 35th is colored purple on both protein alignments).

We sought to determine whether the 12 descendent species without a clear candidate HIS-35 gene represent true losses or merely failures of gene annotation. TBLASTN and BLASTN searches failed to recover likely HIS-35 orthologs absent from the annotation. In contrast to the case of HTAS-1 above, we are confident in the ability of these methods to detect HIS-35 orthologs, since we were able to detect the annotated HIS-35 ortholog by identical methods (i.e., the positive control). For instance, using *C. japonica* HIS-35 as a query, BLASTN searches revealed the (annotated) putative ortholog in *C. macrosperma*, but did not reveal unannotated candidate orthologs in *C. nouroguensis, C. waitukubuli, C. panamensis*, or *C. becei*, which are equally closely related to *C. japonica*. In addition, whereas species lacking HTAS-1 appear to be randomly scattered across the HTAS-1-containing clade, as expected by random gene annotation failures, species lacking HIS-35 within the HIS-35-containing clade group in subclades. This is as expected by true biological loss and not by random annotation failures. Thus, we conclude that HIS-35 has been lost some 5 times independently in different *Caenorhabditis* lineages.

### Conservative protein evolution of HIS-35 and evidence for occasional gene conversion with core H2A

HIS-35 provides a particularly interesting example for histone protein evolution. In *C. elegans*, the protein sequence of HIS-35 differs by just one amino acid from the S-phase H2A despite the substantial evolutionary time. Namely, HIS-35 has an “A”, while H2A has a “G” at position 124 of the amino acid sequence. If an “A” at this position is an overriding change in the HIS-35 variant, then we could expect this change to encode a different function from the canonical H2A.

However, when we looked at position 124 the canonical H2A sequences of all the *Caenorhabditis* species we actually found out that the “A” is ancestral to canonical H2A (Figure 3). This conservation of the “A” at position 124 suggests that HIS-35 likely has not diverged in function from the ancestral H2A despite this variant being retained across many *Caenorhabditis* species. We also found that the predicted protein sequences of HIS-35 and H2A of species *C. kamaaina* are exactly the same (Figure 3). These findings are not as expected if canonical H2A and HIS-35 proteins have different functions, but are as expected if the proteins are functionally redundant.

The multiple copies of core histone genes are known to undergo so-called concerted evolution, with sequences being transferred between gene copies by recombination (72-75). We, therefore, wondered whether concerted evolution plays a role in the identical protein sequence changes observed in the H2A and HIS-35 paralogs of some species. Under concerted evolution, the two sequences undergoing concerted evolution are homogenized (one overwrites the other). Consequently, the prediction is that such events should lead the interconverting partners to the group together on a phylogenetic tree. To search for evidence of concerted evolution, we reconstructed separate phylogenetic trees of exon-1 and exon-2 for all H2A and HIS-35 sequences. Most of the reconstructed tree largely reflected the species tree, suggesting against the possibility of widespread concerted evolution (Figures S3 and S4). However, we did observe the grouping of the two gene sequences for *Caenorhabditis sp21*, consistent with the concerted evolution of HIS-35 and H2A in only this species. Concerted evolution of these genes is consistent with a lack of functional differentiation, though admittedly the low rate of such events weakens the strength of this argument.

### The intron loss and gain in HTZ-1 show a dynamic evolutionary history

The ubiquitously expressed *C. elegans* H2A variant H2A.Z^HTZ-1^ is the ortholog of H2A.Z, which is evolutionary conserved across all the eukaryotes (3, 76). *C. elegans* H2A.Z^HTZ-1^ has an intron that splits the 57th codon at position 2 (Figure 1). Within the alignment across all H2A variants, we searched for genes that share the single intron position of *C. elegans* H2A.Z^HTZ-1^, revealing 30 genes share this position (Figure 4, marked with a plus sign; Figure S1). These are putative H2A.Z^HTZ-1^ orthologs; consistent with this hypothesis, they grouped together on the tree. Unexpectedly, we found that 22 of these 30 putative H2A.Z^HTZ-1^ genes have two (or more in *Diploscapter*) introns in their genes, one at position 57.2 and the other at position 111.1. Both introns have been repeatedly individually lost in different lineages (including in the lineage leading to the single-intron *C. elegans* H2A.Z^HTZ-1^ gene). Species including *C. elegans, C. tropicallis, C. sp32, C. afra, C. guadeloupensis, C. virilis* have lost the second H2A.Z intron which is at position 111.1, whereas *C. casteli* and *C. angaria* have lost their first intron. These results are consistent with general results in protein-coding genes, wherein intron loss is common across the *Caenorhabditis* phylogeny (77, 78). Nonetheless, the finding that general trends of intron loss may equally apply to histone variant genes is important in understanding the functional implications of variant introns.

**Figure 4:**
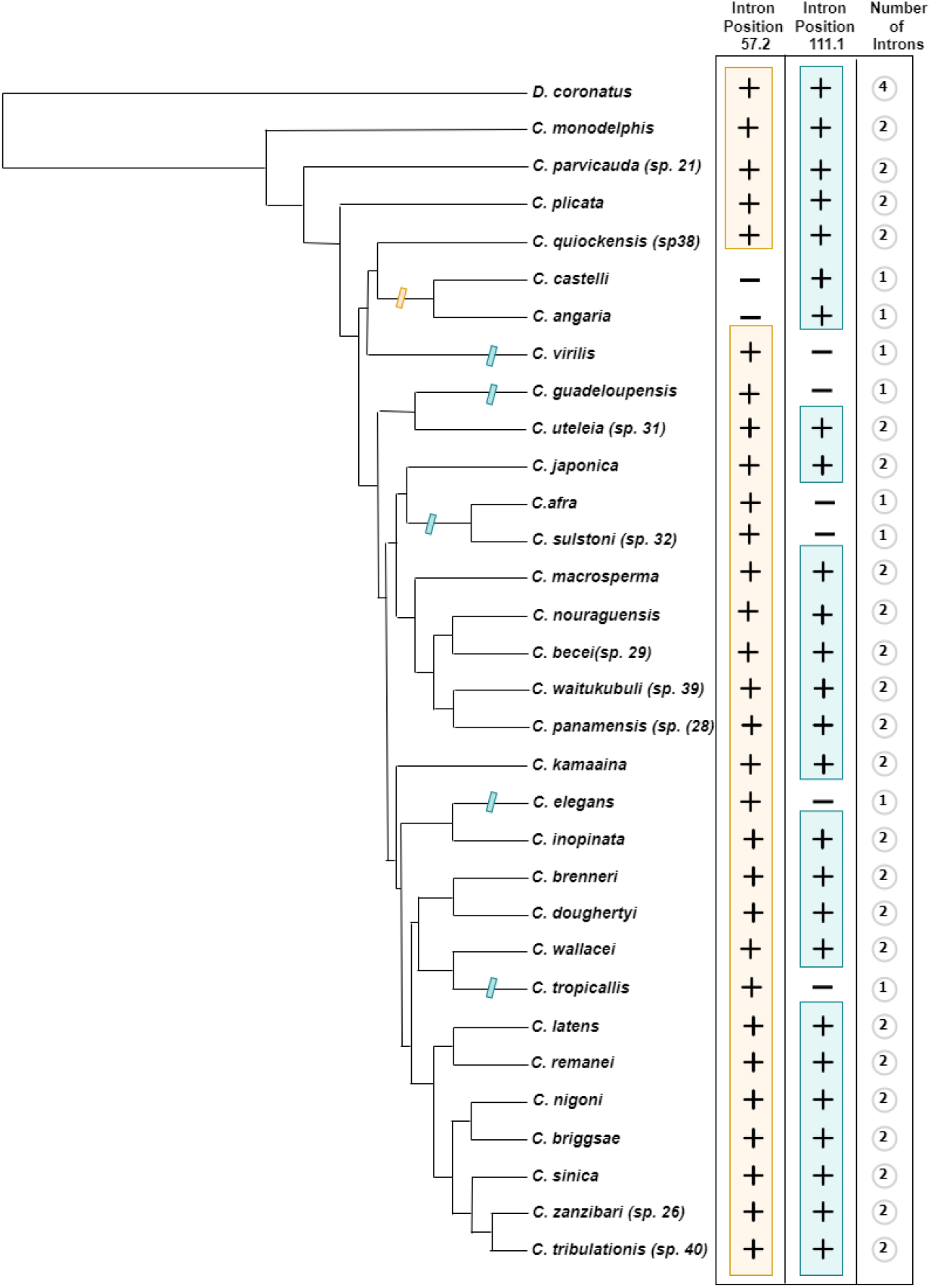
The dynamic history of intron loss and gain in HTZ-1. On the left is the previously reconstructed species tree of *Caenorhabditis*. HTZ-1 characteristic intron presence and absence in a species is indicated by the plus and minus sign. Yellow hash marks on the tree branch depict the loss of intron-1 whereas the blue hash mark on the branch suggests the loss of intron 2 in those lineages.

Intron presence is a conspicuous difference between core and variant histones, raising the question of the functional significance of variant histone introns: specifically, are variant histone introns important for differentiating the specific functions of those variants from their core paralogs? The finding of recurrent loss of introns from a variant gene suggests that the specific positions or sequences of introns in a particular histone variant gene may not be of particular importance. Nonetheless, it is of note that all observed H2A.Z orthologs contain at least one intron, suggesting that the presence of at least one intron, at whatever position, could be important for efficient expression of variant histone genes, consistent with their expression through canonical gene expression pathways, in which intron presence often promotes gene expression.

## Discussion

### Introns as sources of phylogenetic information

This study is among the first to leverage the information present in intron positions to decipher the evolutionary history of histone variants (60). Previous studies have shown intron position conservation among widely diverged eukaryotic species (53-55, 57-60). For instance, intron positions are highly conserved between humans, mice, and fish (57). Thus, intron positions contain a record of evolutionary history that can facilitate insights into gene history. The utility of introns here is different for the different variants and allows different questions to be addressed. The clearest case comes from HIS-35, in which nearly complete evolutionary conservation of H2A and HIS-35 sequences leads to a lack of phylogenetic signal within proteins. At the other end of the spectrum lies HTAS-1, which shows much more rapid sequence evolution. However, here, the discrepancy between rates between HTAS-1 and H2A, and likely between different sites in the two proteins (i.e., the N and C termini are highly constrained in H2A but fast-evolving for HTAS-1) makes it impossible to define a single evolutionary model across the gene family. As expected by general long branch attraction consideration, this leads to fast-evolving HTAS-1 incorrectly grouping far away from *Caenorhabditis* core H2A sequences. This generally undermines our confidence in sequence-based phylogenetic reconstruction of HTAS-1 genes. Ironically, intron-defined HTAS-1 orthologs do group as a clade, indicating that sequence-based phylogenetic reconstruction was likely successful for this group; however, without the intron position information, our general lack of confidence in the methods’ success for HTAS-1 would lead us to question this finding. Thus, for this case, even though the sequence-based phylogenetic methods apparently correctly identified the HTAS-1 clade, the orthologous information from intron positions was necessary for us to be confident in the sequence-based phylogeny. Importantly, intron positions were indispensable in distinguishing the origins of HTAS-1. Because HTAS-1 arose in an ancestor containing two genes encoding nearly identical proteins (H2A and HIS-35), it is very difficult to determine whether H2A evolved from the preexisting variant HIS-35 or *de novo* from H2A. The fact that all candidate HTAS-1 genes lack the HIS-35 intron strongly suggests *de novo* evolution from H2A, not HIS-35.

### Evidence from phylogenetic analysis for functional significance of recently-evolved histone variants

In addition to tracing the origins and subsequent history of gene loss and retention, our results provide insights into the possible functions of histone H2A variants. For example, HIS-35 differs by a single amino acid from the S-phase H2A (‘A’ instead of ‘G’ at the 124th position). Given that characterized histone variants are thought to largely represent functionally distinct proteins, one hypothesis is that this single difference functionally differentiates HIS-35 protein from H2A. One way that a single amino acid change could have outsize effects is through altering the landscape of posttranslational modifications, which are key to histone function. For instance, this is the case with H3 variants, H3.1 and H3.2 which differ from one another by single amino acid and show distinct patterns of expression and post-translational modifications (64, 65).

Given it is possible for a small amino acid difference to change function, we approached the single H2A/HIS-35 difference with the hypothesis that the single difference could lead to functionally different proteins. However, when we looked at the H2A sequences of all the *Caenorhabditis* species, we found an ‘A’ at position 124 to be ancestral. Considering the presence of an A at position 124 in other canonical H2As, suggests the variant HIS-35 might have the same function as the canonical H2A. Lack of functional differentiation is also consistent with the similarity of A and G amino acids. This hypothesis is also supported by the case of *C. kamaanina*, in which the encoded HIS-35 and H2A protein sequences are exactly the same. While these results do not disprove the hypothesis that H2A and HIS-35 encode proteins with important functional differences (except in the case of *C. kamaanina*), we propose instead that HIS-35 protein’s functional importance lies in allowing for the expression of a protein with overlapping or redundant functions to canonical H2A that is not restricted to S-phase, as is the case with canonical H2A. This could allow expression in differentiated cells that do not undergo mitosis and enable tissue-specific expression. Such a potential semi-redundancy could help to explain the ambivalent phylogenetic pattern, in which retention of HIS-35 in most species suggests functional importance whereas loss in 5 independent lineages suggests conditional expendability. Interestingly, a similar pattern of lineage-specific loss has been observed for H2B variants, which has also been interpreted as encoding functionally important but partially redundant functions (79).

### Function in reproduction for the sperm-specific variant HTAS-1

Sperm-specific proteins show generally elevated rates of evolution, consistent with strong selection on sperm functions because of sperm competition (67, 80). The current data show that this is decisively the case for the sperm-specific variant HTAS-1. We show that *C. elegans* HTAS-1’s greater divergence from core H2A is not because of differences in an evolutionary age, but very much despite it: HTAS-1 is most divergent variant protein despite being the most recent to diverge from core H2A, having since evolved at a rate many times higher than any other H2A paralog. This high rate of evolution strongly suggests that HTAS-1 may be adapted to play roles important for reproduction in some species.

However, our data also have somewhat ambiguous implications for HTAS-1 function. On the one hand, though HTAS-1 has been maintained for long periods of time in many lineages it may have also been lost in multiple independent lineages, as found for HIS-35. While rapid evolution of HTAS-1 made it impossible to exclude the possibility that apparent HTAS-1 losses actually represent gene annotation failures, the fact that careful scrutiny revealed no gene annotation failures for HIS-35 weighs against the possibility of annotation failure, suggesting real loss of HTAS-1.

On the other hand, the much larger degree of protein sequence difference between HTAS-1 and H2A would seem to decrease the probability that HTAS-1 protein is functionally identical to H2A protein particularly given the extended C & N terminus of HTAS-1 which has previously been reported to play a vital role in DNA compaction, chromosome segregation, and fertility (67). Moreover, the particular chromatin constraints of sperm production raise the possibility that HTAS-1 proteins could encode distinct functions relative to H2A proteins, for instance by leading to greater sperm DNA compaction; however, it is also possible that a distinct H2A paralog is simply necessary to ensure expression of H2A proteins well after germline mitosis in later stages of spermatogenesis that undergo transcription, DNA recombination and repair, and division. Thus, more study of the functional significance of HTAS-1 homologs in different species is clearly needed to distinguish between these possibilities.

### Concluding remarks

To summarize, these results show exceptions to previously reported patterns, challenging sometimes implicit assumptions about non-core histones. First, whereas protein sequence differences between core and variant histone paralogs are often assumed to reflect differences in protein function, here we show that the variant protein HIS-35 is likely to have a redundant function with core H2A despite the sequence difference. Second, while all *C. elegans* H2A variants have a single intron, our observation of multi-intron variants and of recurrent intron loss, suggests that specific introns may not have crucial roles in the expression of histone variants. Instead, the role of introns in variant histones may simply lie in introns’ general roles in promoting expression. Third, the combination of conservation and loss of variant histones points to potentially lineage-specific, partially redundant, or easily replaced roles of some histone variants. Future studies should explore the generality of these patterns across other lineages of eukaryotes. In addition to our specific findings for histone variant biology, these results highlight that introns can be useful in the reconstruction of the histories of complex gene families.

## MATERIAL AND METHODS

### Data source

Genomic sequences and gene feature format files of 108 nematode species were obtained from WormBase and the *Caenorhabditis* database (68, 69).

### Data mining and processing

All the known genes of 108 nematode species with characterized exon-intron structures were fetched from their genomes using their respective ‘gene feature format’ file. We then noted the positions of the introns in the header of their respective genes and translated the *Caenorhabditis* gene sequences.

To identify the homologs of H2A and their variants, BLASTP, version 2.9.0+, was performed using standard parameters while treating the translated gene sequences (of 108 nematode species) as the database and H2A and variant (H2A.Z^HTZ-1^, HTAS-1, HIS-35) protein sequences as the query (81). Using a maximum e-value of 1e-10, 8003 hits were retrieved which were the homologs of H2A and H2A-variant genes. We then removed dubious genes encoding proteins more than 200 amino acids long, because histone proteins are generally shorter. We collapsed the genes whose introns align at the position and have an identical protein sequence. After filtering off the genes we were left with 408 distinct entries.

Previous studies have shown the intron position conservation among widely diverged eukaryotic species (53-55, 57-60). Therefore, to assess the intron position conservation among the putative H2A variant genes, we performed a Multiple Sequence Alignment (MSA) using the default parameters of CLUSTALW (82). We mapped the intron positions of each gene onto the corresponding protein CLUSTALW alignment, allowing us to identify as potential H2A.Z^HTZ-1^, HTAS-1, HIS-35 orthologs those genes with intron positions matching *C. elegans* intron positions.

### Phylogenetic Analysis

Multiple sequence alignment of 408 H2A variants homologs was performed using default parameters of MUSCLE to generate a phylogenetic tree (Figure S1) using IQtree, which does an automatic selection of the model by doing a model fit test and likelihood scoring (83, 84). VT+R9, the variable time method, was selected by IQtree for our data. The tree didn’t yield a clear phylogenetic signal for HIS-35 or H2A.Z^HTZ-1^, with homologs exhibiting the *C. elegans* HIS-35 or H2A.Z^HTZ-1^ intron positions are scattered over the tree (Figure S1; IP 234 and IP 233 respectively). However, when we took a closer look at the HIS-35 characteristic intron-containing genes, we see that genes containing an intron at the *C. elegans* HIS-35 intron position (Figure S1, IP 234) are restricted to most species of *Caenorhabditis* and its sister genus *Diploscapter*. A clear clade of species was seen which had HTAS-1 characteristic intron position (Figure S1, IP 152). For consistency in Figures 2-4, we show the portion of the phylogenetic tree containing *Caenorhabditis* species and *Disploscapter*.

### Confirmation of H2A variant losses

We found a loss of H2A.Z^HTZ-1^, HTAS-1, HIS-35 characteristic introns in a few lineages (marked by a minus sign in Figures 2, 3, and 4). To know whether this is a real loss or reflected errors in gene annotation, tblastn searches were performed across the genome of these species. This manual curation led to the variant’s characteristic intron splice sites being identified by eye in a few species due to alignment gaps at the exact intron position, indicating that these species truly contain the variant and that failure to initially identify the variant is due to a failure of the annotation to include these genes. We included a material and method flowchart as the Figure S4.

## Acknowledgments

We thank our lab members for their helpful discussions. This work was supported by NSF:1616878; NSF STC DBI-1548297; NSF MCB RUI 1817611; NIH NICHD R03HD093990.

