## Supplementary Figures for "Nematode histone H2A variant evolution reveals diverse histories of retention and loss and evidence for conserved core-like variants"

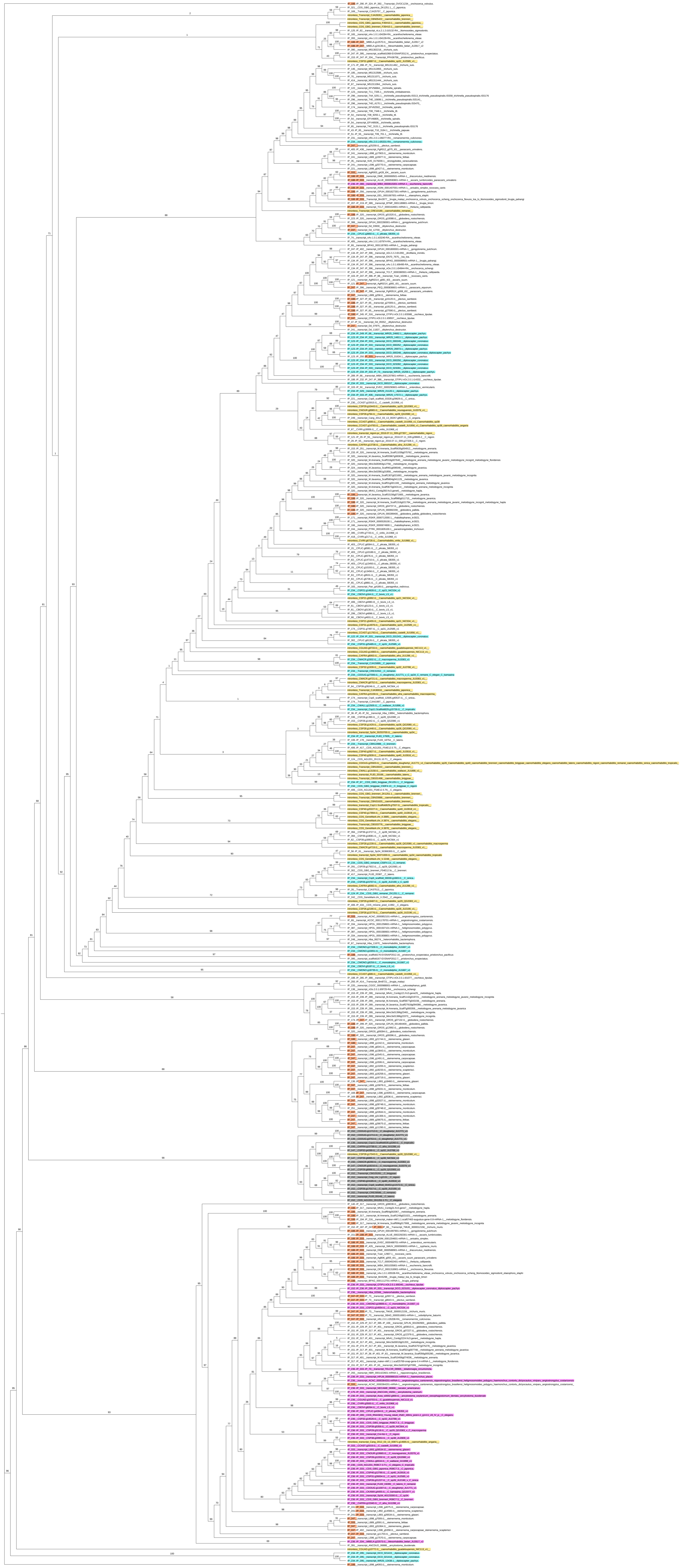

**Figure S1**

Phylogenetic tree reconstructed from protein sequences encoded by H2A and H2A-related variant genes across 108 available nematode genomes. The tree is made up of 408 unique protein entries, with genes encoding identical protein sequences and with identical intron positions collapsed to a single entry (in which cases all collapsed species are indicated). On the tree, "IP" followed by a number stands for the position at which the intron was aligned during the multiple sequence alignment. The intron position of *C. elegans* HTAS-1 is 152 (IP 152), highlighted in grey. The intron position of *C. elegans* HIS-35 is 234 (IP 234), highlighted in blue and the intron position of the *C. elegans* HTZ-1 is 236 (IP 236) highlighted in pink. We have also highlighted the intron positions 188, 247, and 333 in orange, considering them to be HTZ-1 since these cases have at least one of the HTZ-1 characteristic intron positions (either 236 or 333). Intronless sequences (most of which are likely canonical H2As) are indicated as "intronless" and they are highlighted with yellow. The intronless H2A copy of *C.Sp 28* is being clustered within the HTAS-1 clade (highlighted in grey) is an incomplete protein sequence.

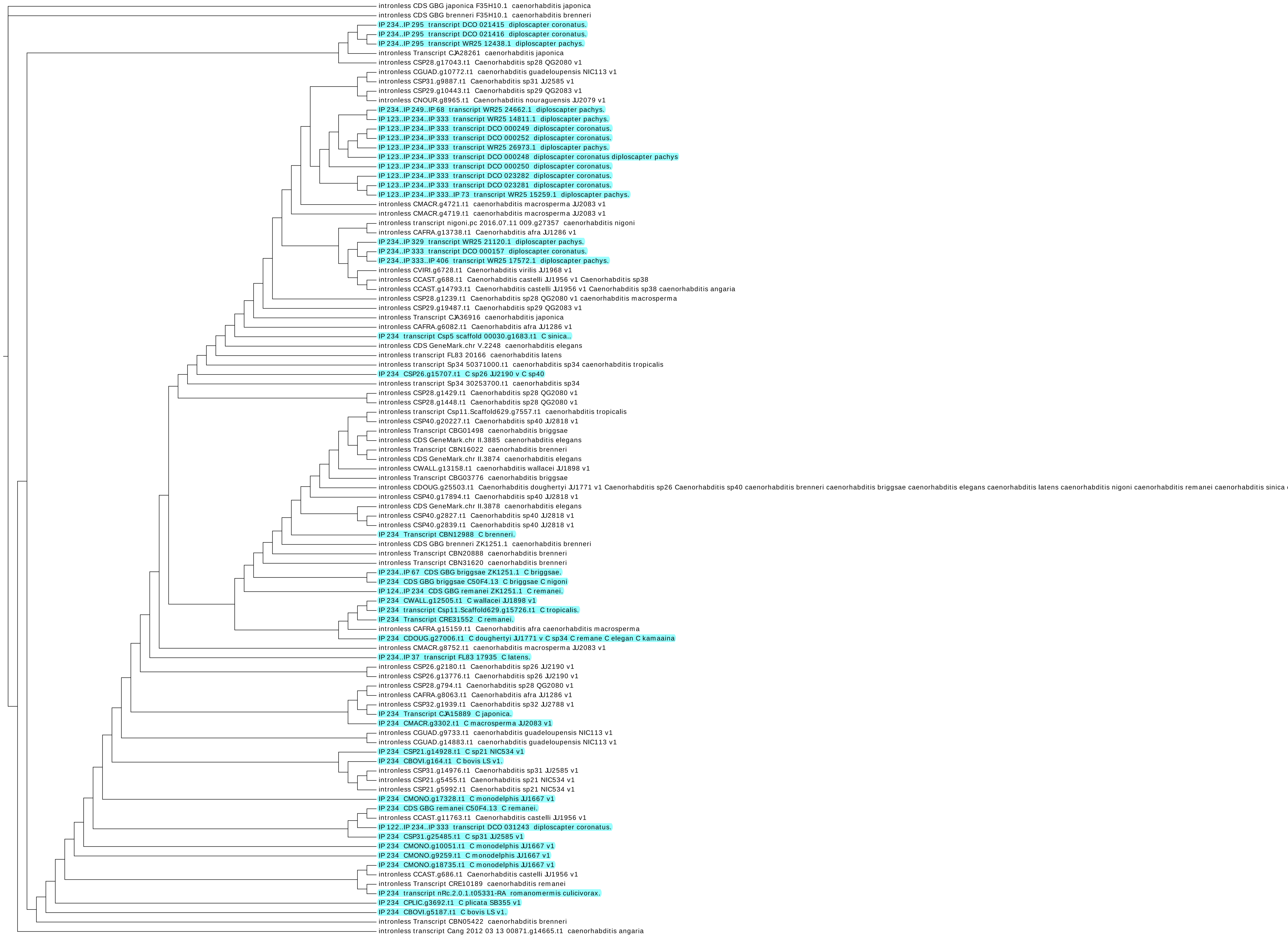
intronless CDS GBG japonica F35H10.1 caenorhabditis japonica
intronless CDS GBG brenneri F35H10.1 caenorhabditis brenneri
IP 234..IP 295 transcript DCO 021415 diploscapter coronatus.
IP 234..IP 295 transcript DCO 021416 diploscapter coronatus.
IP 234..IP 295 transcript WR25 12438.1 diploscapter pachys.
intronless Transcript CJA28261 caenorhabditis japonica
intronless CSP28.g17043.t1 Caenorhabditis sp28 QG2080 v1
intronless CGUAD.g10772.t1 caenorhabditis guadeloupensis NIC113 v1
intronless CSP31.g9887.t1 Caenorhabditis sp31 JU2585 v1
intronless CSP29.g10443.t1 Caenorhabditis sp29 QG2083 v1
intronless CNOUR.g8965.t1 Caenorhabditis nouraguensis JU2079 v1
IP 234..IP 249..IP 68 transcript WR25 24662.1 diploscapter pachys.
IP 123..IP 234..IP 333 transcript WR25 14811.1 diploscapter pachys.
IP 123..IP 234..IP 333 transcript DCO 000249 diploscapter coronatus.
IP 123..IP 234..IP 333 transcript DCO 000252 diploscapter coronatus.
IP 123..IP 234..IP 333 transcript WR25 26973.1 diploscapter pachys.
IP 123..IP 234..IP 333 transcript DCO 000248 diploscapter coronatus diploscapter pachys
IP 123..IP 234..IP 333 transcript DCO 000250 diploscapter coronatus.
IP 123..IP 234..IP 333 transcript DCO 023282 diploscapter coronatus.
IP 123..IP 234..IP 333 transcript DCO 023281 diploscapter coronatus.
IP 123..IP 234..IP 333..IP 73 transcript WR25 15259.1 diploscapter pachys.
intronless CMACR.g4721.t1 caenorhabditis macrosperma JU2083 v1
intronless CMACR.g4719.t1 caenorhabditis macrosperma JU2083 v1
intronless transcript nigoni.pc 2016.07.11 009.g27357 caenorhabditis nigoni
intronless CAFRA.g13738.t1 Caenorhabditis afra JU1286 v1
IP 234..IP 329 transcript WR25 21120.1 diploscapter pachys.
IP 234..IP 333 transcript DCO 000157 diploscapter coronatus.
IP 234..IP 333..IP 406 transcript WR25 17572.1 diploscapter pachys.
intronless CVIRI.g6728.t1 Caenorhabditis virilis JU1968 v1
intronless CCAST.g688.t1 Caenorhabditis castelli JU1956 v1 Caenorhabditis sp38
intronless CCAST.g14793.t1 Caenorhabditis castelli JU1956 v1 Caenorhabditis sp38 caenorhabditis angaria
intronless CSP28.g1239.t1 Caenorhabditis sp28 QG2080 v1 caenorhabditis macrosperma
intronless CSP29.g19487.t1 Caenorhabditis sp29 QG2083 v1
intronless Transcript CJA36916 caenorhabditis japonica
intronless CAFRA.g6082.t1 Caenorhabditis afra JU1286 v1
IP 234 transcript Csp5 scaffold 00030.g1683.t1 C sinica..
intronless CDS GeneMark.chr V.2248 caenorhabditis elegans
intronless transcript FL83 20166 caenorhabditis latens
intronless transcript Sp34 50371000.t1 caenorhabditis sp34 caenorhabditis tropicalis
IP 234 CSP26.g15707.t1 C sp26 JU2190 v C sp40
intronless transcript Sp34 30253700.t1 caenorhabditis sp34
intronless CSP28.g1429.t1 Caenorhabditis sp28 QG2080 v1
intronless CSP28.g1448.t1 Caenorhabditis sp28 QG2080 v1
intronless transcript Csp11.Scaffold629.g7557.t1 caenorhabditis tropicalis
intronless CSP40.g20227.t1 Caenorhabditis sp40 JU2818 v1
intronless Transcript CBG01498 caenorhabditis briggsae
intronless CDS GeneMark.chr II.3885 caenorhabditis elegans
intronless Transcript CBN16022 caenorhabditis brenneri
intronless CDS GeneMark.chr II.3874 caenorhabditis elegans
intronless CWALL.g13158.t1 caenorhabditis wallacei JU1898 v1
intronless Transcript CBG03776 caenorhabditis briggsae
intronless CDOUG.g25503.t1 Caenorhabditis doughertyi JU1771 v1 Caenorhabditis sp26 Caenorhabditis sp40 caenorhabditis brenneri caenorhabditis briggsae caenorhabditis elegans caenorhabditis latens caenorhabditis nigoni caenorhabditis remanei caenorhabditis sinica
intronless CSP40.g17894.t1 Caenorhabditis sp40 JU2818 v1
intronless CDS GeneMark.chr II.3878 caenorhabditis elegans
intronless CSP40.g2827.t1 Caenorhabditis sp40 JU2818 v1
intronless CSP40.g2839.t1 Caenorhabditis sp40 JU2818 v1
IP 234 Transcript CBN12988 C brenneri.
intronless CDS GBG brenneri ZK1251.1 caenorhabditis brenneri
intronless Transcript CBN20888 caenorhabditis brenneri
intronless Transcript CBN31620 caenorhabditis brenneri
IP 234..IP 67 CDS GBG briggsae ZK1251.1 C briggsae.
IP 234 CDS GBG briggsae C50F4.13 C briggsae C nigoni
IP 124..IP 234 CDS GBG remanei ZK1251.1 C remanei.
IP 234 CWALL.g12505.t1 C wallacei JU1898 v1
IP 234 transcript Csp11.Scaffold629.g15726.t1 C tropicalis.
IP 234 Transcript CRE31552 C remanei.
intronless CAFRA.g15159.t1 Caenorhabditis afra caenorhabditis macrosperma
IP 234 CDOUG.g27006.t1 C doughertyi JU1771 v C sp34 C remane C elegan C kamaaina
intronless CMACR.g8752.t1 caenorhabditis macrosperma JU2083 v1
IP 234..IP 37 transcript FL83 17935 C latens.
intronless CSP26.g2180.t1 Caenorhabditis sp26 JU2190 v1
intronless CSP26.g13776.t1 Caenorhabditis sp26 JU2190 v1
intronless CSP28.g794.t1 Caenorhabditis sp28 QG2080 v1
intronless CAFRA.g8063.t1 Caenorhabditis afra JU1286 v1
intronless CSP32.g1939.t1 Caenorhabditis sp32 JU2788 v1
IP 234 Transcript CJA15889 C japonica.
IP 234 CMACR.g3302.t1 C macrosperma JU2083 v1
intronless CGUAD.g9733.t1 caenorhabditis guadeloupensis NIC113 v1
intronless CGUAD.g14883.t1 caenorhabditis guadeloupensis NIC113 v1
IP 234 CSP21.g14928.t1 C sp21 NIC534 v1
IP 234 CBOVI.g164.t1 C bovis LS v1.
intronless CSP31.g14976.t1 Caenorhabditis sp31 JU2585 v1
intronless CSP21.g5455.t1 Caenorhabditis sp21 NIC534 v1
intronless CSP21.g5992.t1 Caenorhabditis sp21 NIC534 v1
IP 234 CMONO.g17328.t1 C monodelphis JU1667 v1
IP 234 CDS GBG remanei C50F4.13 C remanei.
intronless CCAST.g11763.t1 Caenorhabditis castelli JU1956 v1
IP 122..IP 234..IP 333 transcript DCO 031243 diploscapter coronatus.
IP 234 CSP31.g25485.t1 C sp31 JU2585 v1
IP 234 CMONO.g10051.t1 C monodelphis JU1667 v1
IP 234 CMONO.g9259.t1 C monodelphis JU1667 v1
IP 234 CMONO.g18735.t1 C monodelphis JU1667 v1
intronless CCAST.g686.t1 Caenorhabditis castelli JU1956 v1
intronless Transcript CRE10189 caenorhabditis remanei
IP 234 transcript nRc.2.0.1.t05331-RA romanomermis culicivorax.
IP 234 CPLIC.g3692.t1 C plicata SB355 v1
IP 234 CBOVI.g5187.t1 C bovis LS v1.
intronless Transcript CBN05422 caenorhabditis brenneri
intronless transcript Cang 2012 03 13 00871.g14665.t1 caenorhabditis angaria

**Figure S2**

Phylogenetic reconstruction on core H2A orthologs and on the HIS-35 orthologs identified based on intron positions. Here, we expect distinct clades of H2A and HIS-35 sequences, yet we recover no such separate clades (Figure S2, HIS-35 is highlighted).

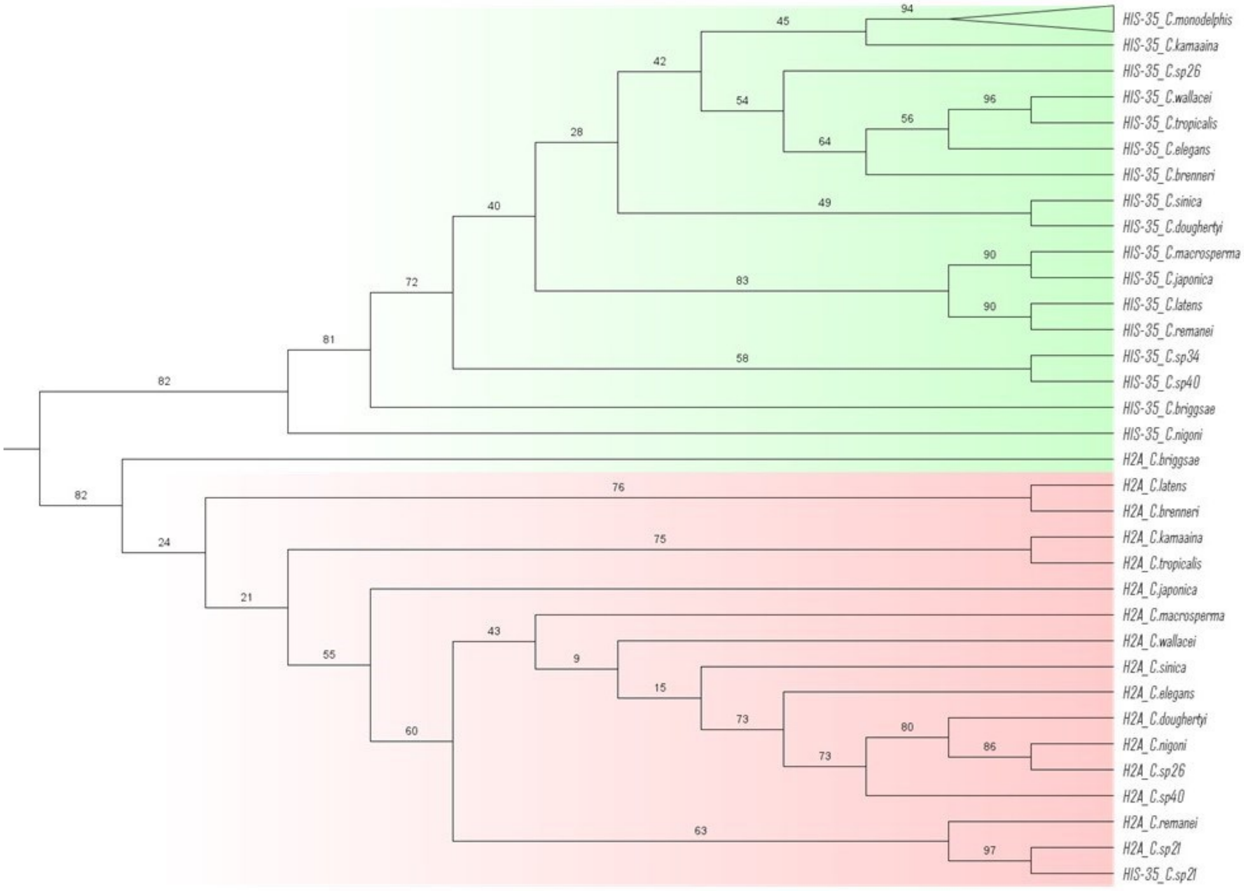


**Figure S3**

Phylogenetic tree reconstructed from nucleotide sequences of canonical *Caenorhabditis* H2A gene sequences and first exons of candidate HIS-35 gene sequences. We see a clear cluster of H2A nucleotide sequences and HIS-35 with a few exceptions, in particular *Caenorhabditis sp 21.*

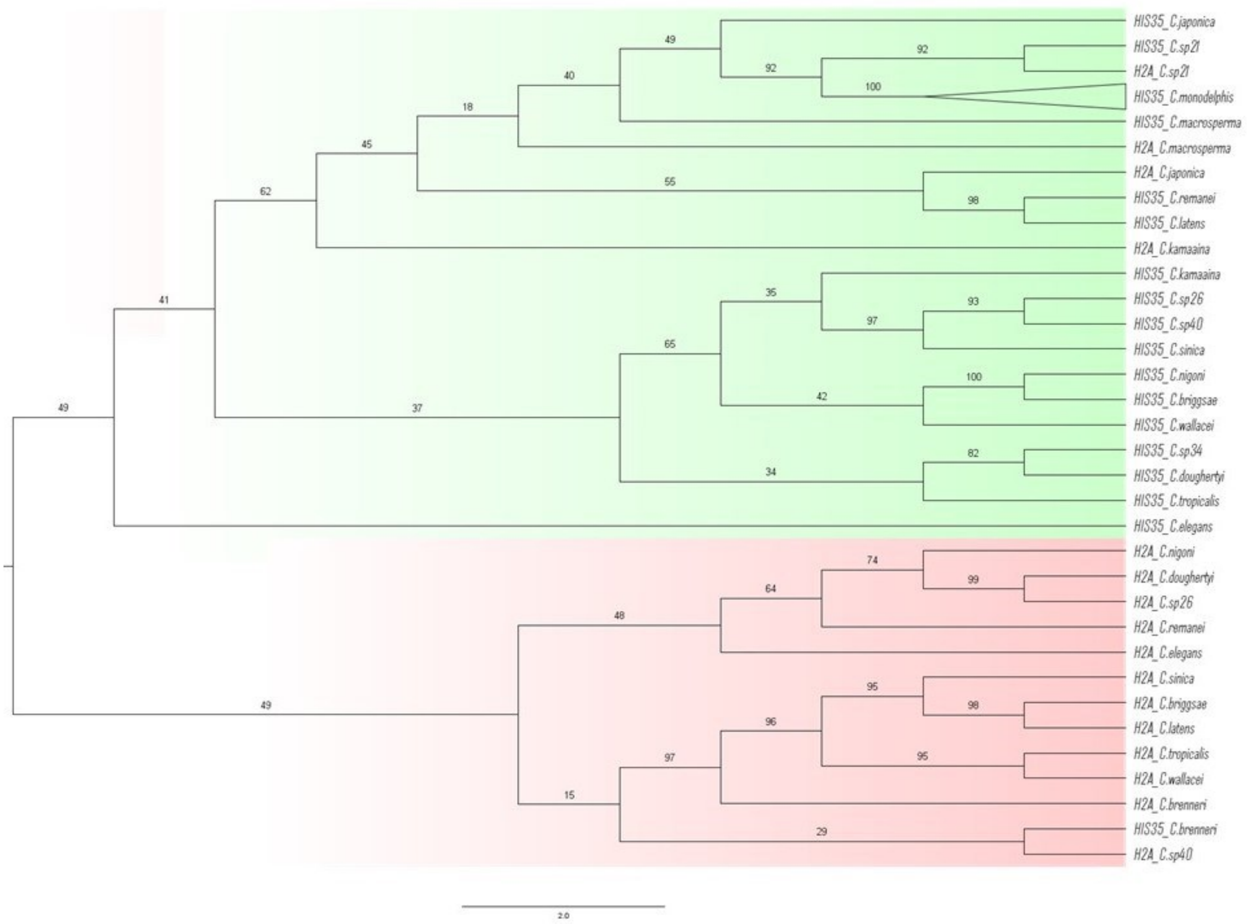


**Figure S4**

Phylogenetic tree reconstructed from nucleotide sequences of canonical *Caenorhabditis* H2A gene sequences and second exons of candidate HIS-35 gene sequences.

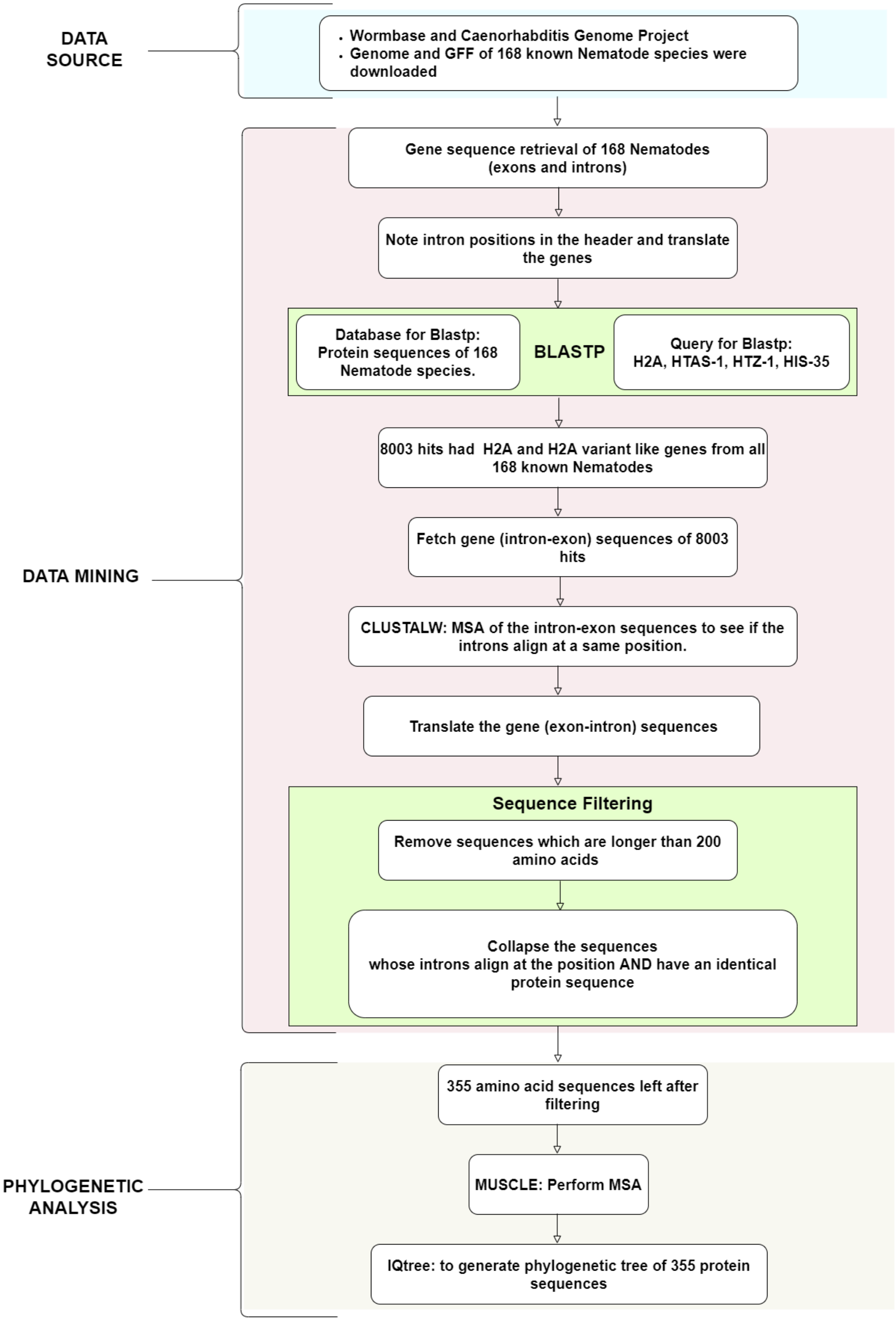


### Figure S5

Material and methods flowchart.
